# Aureochromes are necessary for maintaining polyunsaturated fatty acid content in *Nannochloropsis oceanica*

**DOI:** 10.1101/2021.01.27.428447

**Authors:** Eric Poliner, Andrea W. U. Busch, Linsey Newton, Young Uk Kim, Rachel Clark, Sofía C. Gonzalez-Martinez, Byeong-ryool Jeong, Beronda L. Montgomery, Eva M. Farré

## Abstract

*Nannochloropsis oceanica*, as other stramenopile microalgae, is rich in long-chain polyunsaturated fatty acids (LC-PUFA) such as eiconsapentaenoic acid (EPA). We observed that fatty acid desaturases (FAD) involved in LC-PUFA biosynthesis were among the strongest blue light induced genes in *N. oceanica* CCMP1779. Blue light was also necessary for maintaining LC-PUFA levels in CCMP1779 cells, and growth under red light led to a reduction in EPA content. Aureochromes are stramenopile specific proteins that contain a light-oxygen-voltage-sensing (LOV) domain that associates with a flavin mononucleotide and is able to sense blue light. These proteins also contain a bZIP DNA binding motif and can act as blue light regulated transcription factors by associating with a E-box like motif, which we found enriched in the promoters of blue light induced genes. We demonstrated that, *in vitro*, two CCMP1779 aureochromes were able to absorb blue light. Moreover, the loss or reduction of any of the three aureochromes led to a decrease in the blue light specific induction of several FADs in CCMP1779. EPA content was also significantly reduced in *NoAureo* 2 and *NoAureo* 4 mutants. Taken together, our results indicate that aureochromes mediate blue light dependent regulation of LC-PUFA content in *N. oceanica* CCMP1779 cells.

## INTRODUCTION

Marine environments are enriched in blue light as longer wavelengths such as red and far-red cannot reach as deep in the water column. In contrast to land plants, the extent of blue and red light regulation of downstream processes is not well understood in eukaryotic algae. Photosynthetic stramenopiles, called ochrophytes, are major components of the marine phytoplankton (Tragin and Vaulot, 2018). Their genomes encode for several blue light photoreceptors, including cryptochrome family proteins as well as aureochromes, an ochrophyte specific protein family. Although it has been demonstrated that cryptochromes influence light dependent processes in stramenopiles (Coesel et al., 2009), the mechanism by which they influence gene expression is still unclear. Aureochromes contain a light-oxygen-voltage-sensing domain (LOV) and a basic leucine zipper domain (bZIP) (Essen et al., 2017). Aureochromes mediate blue light dependent photomorphogenesis in the Zanthophyceae *Vaucheria frigida* (Takahashi et al., 2007), and are involved in photoacclimation, cell division and blue light dependent gene expression changes in diatoms (Huysman et al., 2013; Schellenberger Costa et al., 2013; Mann et al., 2017; Mann et al., 2020). As do other bZIP transcription factors, aureochromes dimerize and are able to bind directly to DNA. In vitro, blue light can enhance dimerization and/or DNA binding (Essen et al., 2017; Kroth et al., 2017).

We investigated light regulated gene expression in the ochrophyte *Nannochloropsis oceanica* CCMP1779. *Nannochloropsis* species are small unicellular alga with a diameter of about 2 μm. The genus is distributed widely in the marine environment but is also found in fresh and brackish waters (Fawley and Fawley, 2007) and several species are being used as models for understanding lipid metabolism (Poliner et al., 2018; Li-Beisson et al., 2019). These algae have several putative blue light photoreceptors but appear to lack red light photoreceptors (Vieler et al., 2012). *Nannochloropsis* genomes encode for one cryptochrome protein family (CPF) protein and three aureochromes (Vieler et al., 2012).

Long chain polyunstaturated fatty acids are linked to the marine environment (Li-Beisson et al., 2019). Marine algae have high level of LC-PUFAs such as eicosapentaenoic acid, EPA (20:5; number of carbons: number of double bonds) and many contain docosahexaenoic acid, DHA (22:6) (Li-Beisson et al., 2019). Stramenopiles such as diatoms and *Nannochloropsis* species are used as source of LC-PUFA for fish food or human consumption (Adarme-Vega et al., 2014). LC-PUFAs are mainly located in plastid membrane lipid pools (Li-Beisson et al., 2019). Light quality and quantity appear to affect fatty acid composition in algae. Higher intensities promote the production of saturated fatty acids due to the increased accumulation of triacylglycerol (TAG), but there is also an increase in polyunsaturated fatty acids in chloroplast membrane lipids, which has been proposed to help protect from oxidative stress (Sukenik et al., 1989; Sukenik et al., 2009; Alboresi et al., 2016; Sayanova et al., 2017). The effect of light quality on lipid composition appears to vary depending on the species and growth conditions (Das et al., 2011; Chen et al., 2013; Chen et al., 2015). Our results demonstrate that the early light response of several fatty acid desaturase (FAD) genes in *N. oceanica* CCMP1779 are mainly affected by blue light and only minimally by red light. Moreover, aureochromes are required for the blue light induction of these genes and the loss or decrease in aureochrome expression affects LC-PUFA levels.

## RESULTS

### Light quality mediated changes in gene expression in *N. oceanica* CCMP1779

To study early light signaling in *N. oceanica* and avoid as much as possible secondary downstream events, we quantified gene expression changes in dark-adapted cells after a short blue or red light pulse (Fig. 1A). In our experiment 10,231 out of 10,461 annotated genes were expressed with an average TPM (Transcripts Per Kilobase Million) > 0.1 in at least one of the three conditions (Supplemental Dataset S1). We selected genes that were significantly (q-values < 0.001) induced at least two-fold or repressed at least 50% with respect to the dark control. In these experiments, which focused on identifying early light responsive genes, blue light had stronger effects on gene expression. More genes were differentially expressed under blue than under red light (Fig. 1b). In addition, fold change in expression was smaller for the red light treated samples. For example, the median fold change of induced genes was 3 for blue light and 2.5 for red light. Moreover, hierarchical cluster analysis showed that gene expression after red light treatment was more similar to the dark control than after blue light treatment (Supplemental Fig. S1). Most (82%) of the genes induced by red light were also induced by blue light. In contrast, only 32% of the blue light induced genes were also induced by red light (Fig. 1c). We made a similar observation for the genes repressed by light. About 37% of the blue light repressed genes were also repressed by red light but 77% of red light repressed genes were also repressed by blue light. These results support the hypothesis that the main gene expression regulation by red light is not wavelength specific and therefore, might not be regulated by a photoreceptor.

**Figure 1.**
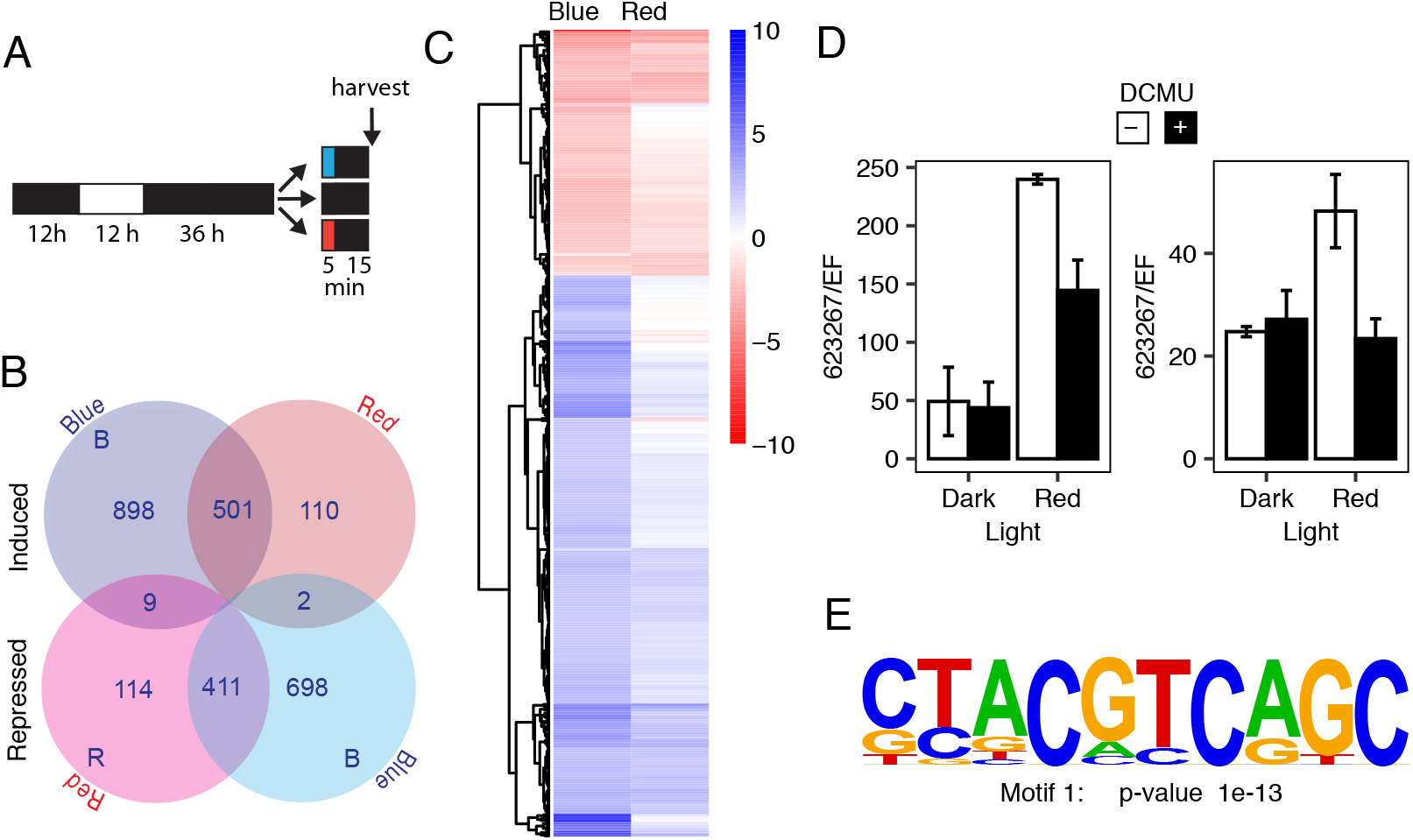
Early light dependent gene expression response in *N. oceanica* CCMP1779. (A) Schematic representation of the experimental set up for the RNA-seq experiment. Cells were grown under light/dark cycles and transferred to the dark for 36 h before being treated 5 minutes light pulse (20 μmol s^−1^ m^−2^). Cells were collected 15 minutes after the end of the light pulse. (B) Number of differentially expressed genes in each category. (C) Hierarchical cluster analysis of differentially expressed genes of their log_2_ transformed fold change in expression with respect to the dark control. (D) DCMU influence on red light induced gene expression. Timing of light treatments as in (A) but red light intensity was 34 μmol s^−1^ m^−2^. Either 20 mM DCMU or ethanol (vehicle control) were added to the cultures 10 minutes before the light pulse. Gene expression was determined by RT-qPCR and shown relative to the elongation factor gene (EF) (average ± range n = 2). Numbers indicate transcript ID. (E) Motif enriched in strongly induced by blue light, identified using HOMER.

To test whether photosynthesis is involved in mediating regulation of gene expression by red light we repeated the light induction experiment in the presence or absence of DCMU (3-(3,4-dichlorophenyl)-1,1-dimethylurea). DCMU inhibits electron transport between photosystem II and plastoquinone, and blocks red light-mediated induction of gene expression in diatoms (Fortunato et al., 2016). We confirmed that this treatment blocked oxygen evolution in *N. oceanica* CCMP1779 (Supplemental Fig. S2). Fig. 2D shows that DCMU reduced the induction of transcripts under red light. These results indicate that an output of photosynthesis mediates regulation of gene expression under these conditions.

**Figure 2.**
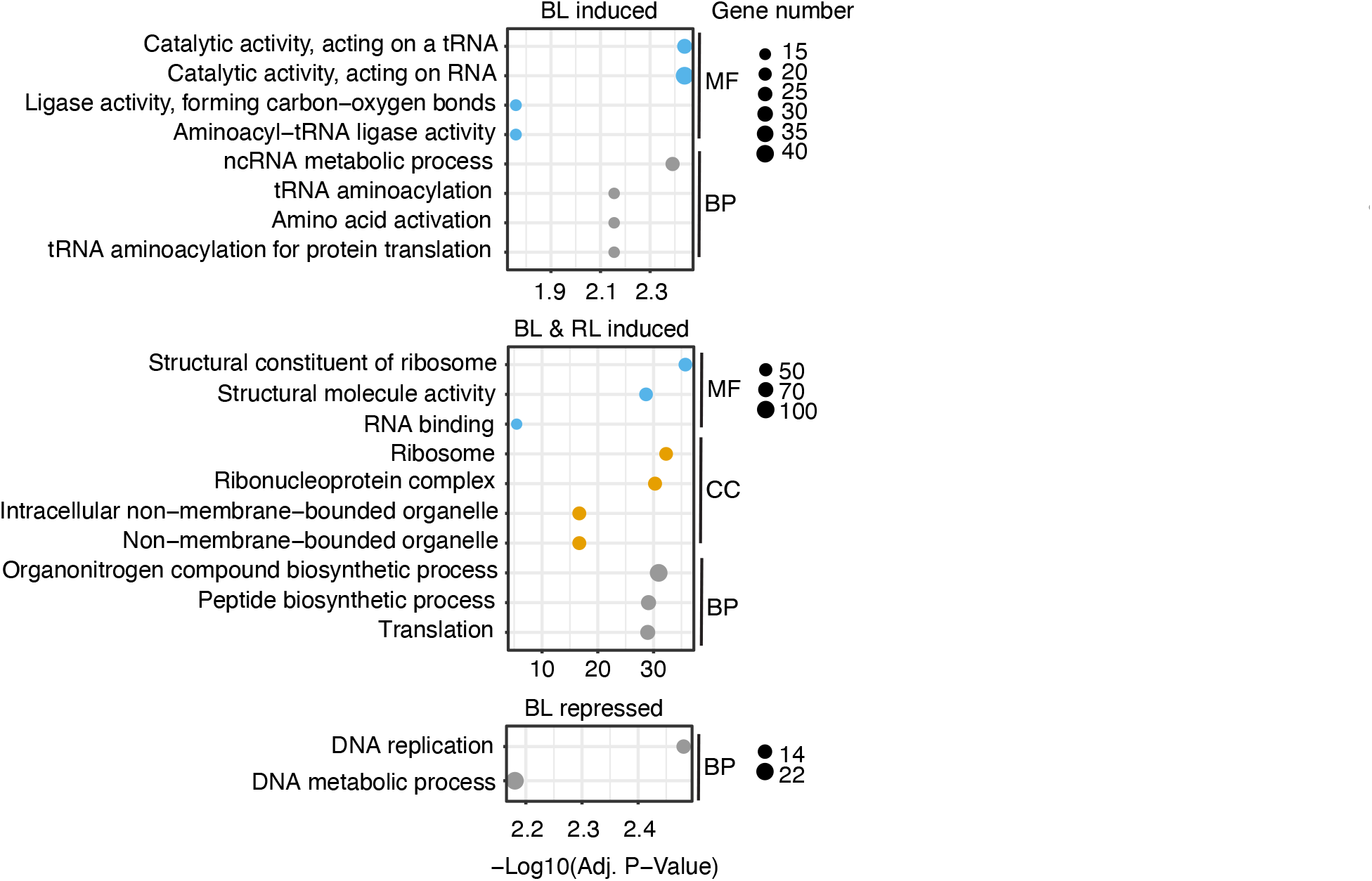
GO enrichment genes of differentially expressed genes. Only categories with significantly enriched functional terms are shown. BL induced, exclusively blue light induced genes; BL & RL induced, genes induced by both blue and red light; RL repressed, genes repressed exclusively by red light. The size of the circles refers to the number of genes in each class. MF, molecular function; BP, biological process; CC, cellular component.

We carried out functional enrichment analyses in gene sets that were either differentially expressed exclusively by blue or red light, and in gene sets induced or repressed by both red and blue light (Fig. 2, Supplemental Fig. S3). No significantly enriched GO terms were found in the sets of genes repressed exclusively by red light or repressed by both red and blue light indicating that a wide range of pathways are affected. Terms related to translation were enriched in blue and blue and red induced genes. For example, exclusively blue light induced genes were enriched in terms related to tRNAs and genes induced by blue and red light were enriched in genes related to ribosomes. The only significantly enriched terms in the blue light repressed genes were related to DNA replication. Analyses using EuKaryotic Orthologous Groups (KOG) led to similar results (Fig. S3). These analyses are limited by the fact that currently only ~50% of genes in *N. oceanica* CCMP1779 contain associated GO (4,795 out of 10,4641) or KOG terms (5,259 out of 10,461).

The genome of *N. oceanica* CCMP1779 encodes for four putative blue light photoreceptors (Vieler et al., 2012). Three of these photoreceptors belong to the aureochrome family. In other stramenopiles, aureochromes bind E-box type DNA elements to regulate gene expression (Takahashi et al., 2007; Heintz and Schlichting, 2016). To test whether aureochromes might be involved in regulating blue light-dependent gene expression in *N. oceanica* CCMP1779 we searched for enriched DNA elements in upstream regions of genes strongly induced by blue but not by red light. The only motif enriched with a p-value < 10^−10^ contained the core sequence of the E-box (ACGT) (Fig. 2E, Supplemental Dataset S2) and is similar to the TGACGT sequence shown to associate with diatom and *Vaucheria* aureochromes (Takahashi et al., 2007; Heintz and Schlichting, 2016). These results indicate that these photoreceptors might play a key role in blue light specific gene expression in *N. oceanica* CCMP1779.

### Blue light induces the expression of fatty acid desaturase genes

Within the group of genes most strongly induced by blue light we identified several fatty acid desaturases (*FAD*) (Fig. 3A). *N. oceanica* CCMP1779 synthesizes EPA (20:5^Δ5, Δ8, Δ11, Δ14, Δ17^) from stearic acid (18:0) via a pathway that involves five desaturation reactions catalyzed by *Δ*9 FAD, *Δ*12 FAD, *Δ*6 FAD, *Δ*5 FAD and *ω*3 FAD (Poliner et al., 2018). The genes encoding for *Δ*9 FAD and *Δ*6 FAD were induced by blue but not by red light. The genes encoding for *Δ*12 FAD, *Δ*5 FAD and *ω3* FAD were strongly induced by blue light but their expression was also elevated, although much less, after the red light treatment. To confirm these results we repeated the experiment and quantified expression using real time quantitative PCR (RT-qPCR) (Fig. 3B). As we observed in our RNA-seq experiment, pulses of blue or white light, induced their expression when measured by RT-qPCR, but red light had a weaker effect. These results indicate that blue light might be necessary for the production of LC-PUFA in *N. oceanica* CCMP1779.

**Figure 3.**
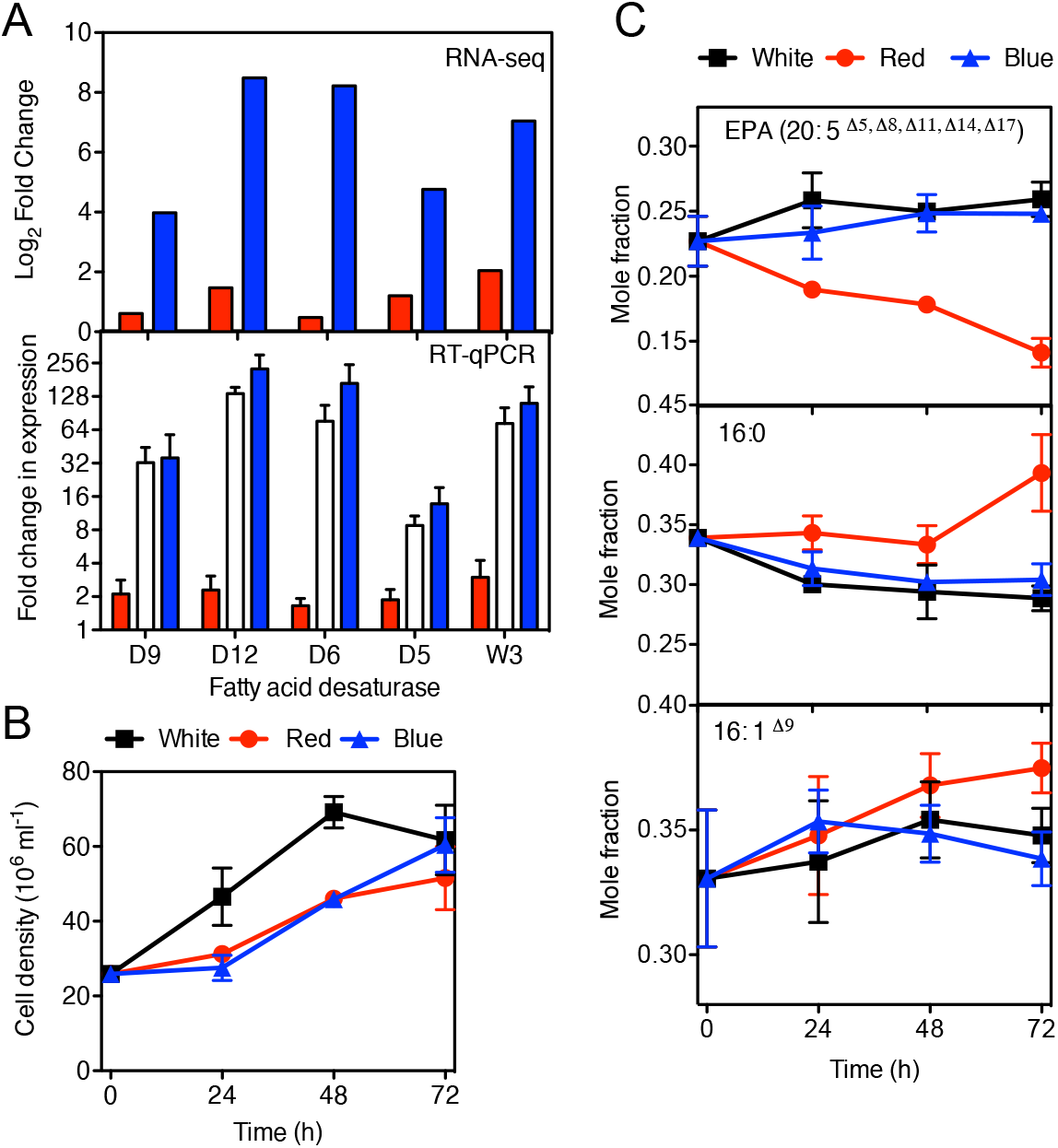
Light quality influences fatty acid desaturase expression and fatty acid composition in *N. oceanica* CCMP1779. (A) Expression of fatty acid desaturase genes after pulses of red, blue or white light. Data from the RNA-seq experiment is shown as log2 transformed fold change in expression with respect to the dark treated samples. A separate experiment measured gene expression changes using RT-qPCR, results are shown as fold change with respect to the average expression of the dark control. The actin-related gene (ACTR) was used as control and the average and standard error are shown (n=3). Growth (B) and fatty acid composition (C) of cells after transfer to either constant white, blue or red light for three days. Cells were grown under constant white light (30 μmol s^−1^ m^−2^), and then transferred to the different light conditions of constant light (30 μmol s^−1^ m^−2^).

To investigate whether fatty acid desaturases are also light regulated in other stramenopiles we searched published diel and light induced gene expression datasets from the diatom *Phaeodactylum tricornutum*. Strong light induction of early light induced genes also occurs at dawn under light/dark cycles (Locke et al., 2005); however, this effect is often masked in diel datasets due to the first light sample being taken several hours after dawn. This was the case for our previous diel *N. oceanica* CCMP1779 dataset, in which the first time point in the light was 3 h after dawn and thus did not enable us to detect acute light responses (Poliner et al., 2015). Acute light responsiveness at dawn should be detectable in experiments in which the first sample in the light was collected 30-60 min after lights on. Of the six putative fatty acid desaturases identified in *P. tricornutum*, five are strongly co-expressed in two available diel studies (Chauton et al., 2013; Smith et al., 2016). Their expression is high during the day and low during the night (Supplemental Fig. S4). These diel experiments were carried out using two different photoperiods, short (8 h light) (Smith et al., 2016), and long (16 h light) (Chauton et al., 2013), and the expression of these genes correlated with these differences. In addition, the data from Chauton et al., which includes timepoints shortly after dawn and dusk, demonstrate that the expression of most of these genes reacts rapidly to light, with an induction just after dawn and a repression at dusk. Finally, all of the six putative fatty acid desaturase genes are induced after a switch from red to blue light in *P. ticornutum* (Mann et al., 2020) and four out of the six are induced by red light in this species (Fortunato et al., 2016) indicating that induction by light is a general feature of fatty acid desaturase genes in stramenopiles with blue light playing a major role.

### Light quality influences fatty acid distribution in *N. oceanica* CCMP1779

To test the potential role of light quality in fatty acid composition we measured fatty acid content in wild type *N. oceanica* CCMP1779 cells grown under white, blue or red light. Cells were grown under constant white light and then either kept in white light or transferred to blue or red of the same light intensities. The growth rate of the cells decreased in blue and red light with respect to white light, but in these experiments the growth rate was similar in both monochromatic lights (Fig. 3B). The mole percentage of EPA decreased in the absence of blue light (Fig. 3C). In contrast, the mole percentage of the unsaturated fatty acid 16:0 and the monounsaturated 16:1^Δ9^ did not decrease in cells grown under red light when compared to cells grown under either blue or white light. These results indicate that blue light is necessary for the maintenance of PUFA content in *N. oceanica* CCMP1779.

### Aureochrome proteins in *N. oceanica* CCMP1779

Since E-box motifs are enriched in the promoter regions of blue light induced genes we investigated whether aureochromes are involved in the blue light mediated signaling that regulates fatty acid desaturation in *N. oceanica* CCMP1779. We cloned and sequenced the coding region of three aureochrome genes using information on *N. oceanica* CCMP1779 v1.0 and v2.0 annotations and aligned sequencing reads (Supplemental Dataset S2). As in other species, *N. oceanica* CCMP1779 aureochromes contain a highly conserved basic region leucine zipper (bZIP) DNA-binding domain, a C-terminal light-oxygen-voltage (LOV) sensing domain and differ in their N and C-terminal extensions (Figure S5A). Phylogenetic analyses using the conserved central bZIP-LOV region indicates that NoAUREO 2 belongs to the Class II (Schellenberger Costa et al., 2013) (Figure S5), NoAUREO3 groups with class I, and NoAUREO 4 groups with Class III/Class IV aureochromes. Some of the apparent differences in N-terminal extensions in the aureochromes of different *Nannochloropsis* species could be caused by differences in gene annotation. For example, in the class I aureochromes, a N-terminal extension is present in *N. salina* CCMP1776 (NSK_000182) and *N. oceanica* CCMP1779 v2.0 annotation available at the Joint Genome Institute (JGI) (jgi|Nanoce1779_2|582424). However, for *N. oceanica* CCMP1779 this extension is not supported by publicly available RNA-seq data (JGI) and is not annotated in *N. gaditana* CCMP526 (Wang et al., 2014).

Interestingly, aureochrome gene expression was influenced by light in our dataset. *NoAUREO* 4 was strongly induced by blue light and *NoAUREO* 2 and *NoAUREO* 3 were slightly, but significantly, repressed by both red and blue light (Supplemental Fig. S6). These results were confirmed in independent experiments (Supplemental Fig. S6).

### NoAUREO3 absorbs blue light

LOV domains of class I aureochromes bind to a flavin chromophore and absorb blue light. When expressed in *E. coli* cells aureochrome 1 from *Vaucheria frigida* and aureochrome 1a from *P. tricornutum*, but not VfAUREO 2 are able to bind a flavin chromophore and absorb blue light (Takahashi et al., 2007; Heintz and Schlichting, 2016). *E. coli* expressing full length NoAUREO 3 had a yellow color indicating the presence of bound flavin (Supplemental Fig. S7A). The expression of full length NoAUREO 2 and NoAUREO 4 was very low in E. coli. NoAUREO 2 expressing cells displayed a slight yellow coloration and NoAUREO 4 expressing cells lacked a visible yellow color (Fig. S7B). Therefore, we focused on the characterization of NoAUREO 3. The absorption spectrum of dark adapted NoAUREO 3 had a maximum at ~450 nm (Figure 4A). Irradiation with blue light led to a decrease in absorption at ~450 nm and an increase in absorption at ~ 392 nm (Fig. 4A). Moreover, when excited at 450 nm the adduct state was not fluorescent but the dark-adapted NoAUREO 3 was (Fig. 4B). At room temperature, the dark reversion kinetics of NoAUREO 3 using absorbance parameters (Fig. 4C) were similar to the ones using fluorescence (Fig. 4D). Our limited analysis of NoAUREO 2 indicated similar fluorescence patterns as NoAUREO 3 (Supplemental Fig. S8), indicating that these *N. oceanica* aureochromes display similar photochemical properties of VfAUREO1 and PtAUREO 1a (Takahashi et al., 2007; Hisatomi et al., 2013; Heintz and Schlichting, 2016).

**Figure 4.**
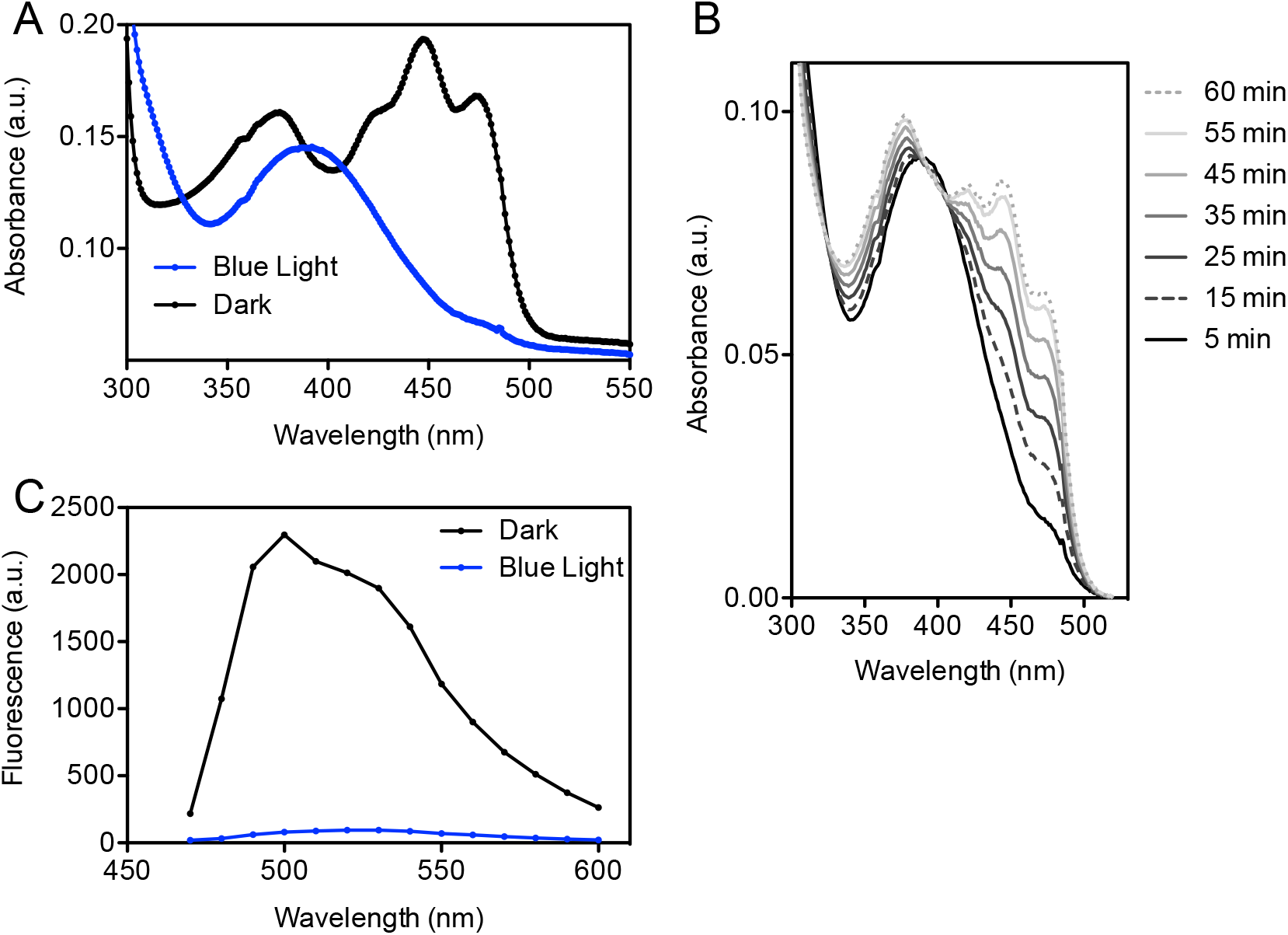
Photochemical characterization of *N. oceanica* CCMP1779 recombinant NoAUREO 3. Absorption spectrum (A) and fluorescence spectrum (B) of recombinant protein in the dark state or after 1 min of blue light treatment. (C) Dark reversion of absorbance spectrum of blue light treated NoAUREO 3, protein was transferred to the dark at time 0. (D) Dark reversion of fluorescence (470-595 nm) of blue light treated NoAUREO 3.

### Generation of Aureochrome mutants

To investigate the role of aureochromes in *N. oceanica* CCMP1779 we generated lines with low or no expression of each of the three aureochromes. Using CRISPR-Cas9 we developed lines that lacked 1542 and 2066 bp of the genomic sequence of *NoAUREO* 2 or *NoAUREO* 4 respectively (Fig. 5A, Supplemental Dataset S2), and therefore were missing most of the coding region. We were unable to isolate *NoAUREO* 3 loss of function mutants using CRISPR-Cas9 in *N. oceanica* CCMP1779. However, the generated *NoAUREO* 3 RNAi lines displayed more than 50% reduction in mRNA content (Fig. 5B).

**Figure 5.**
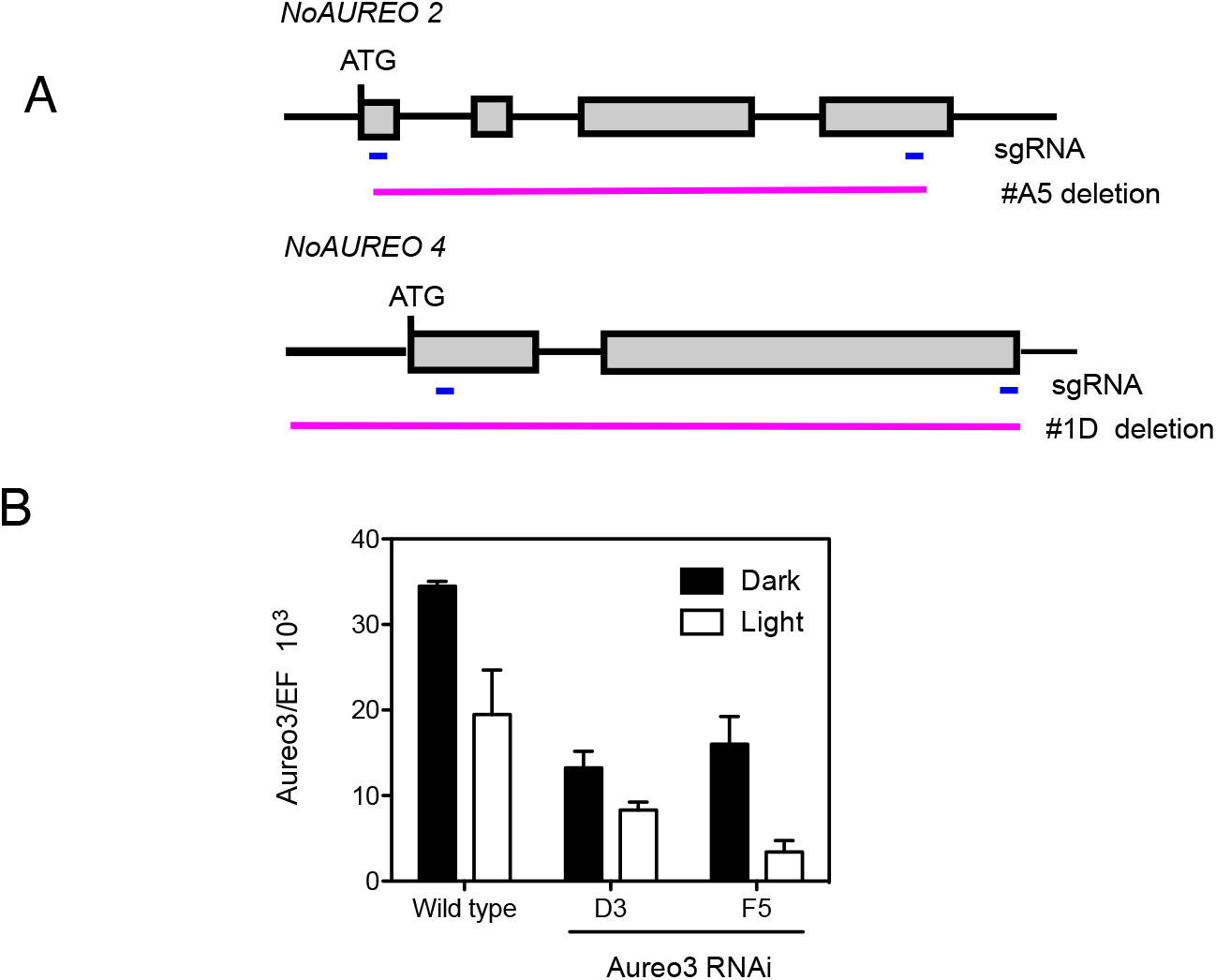
Generation of aureochrome mutants in *N. oceanica* CCMP1779. (A) Schematic representation of Aureo 2-KO and Aureo 4-KO lines. The blue lines indicate the position of the sgRNA and the magenta region the fragment lost in the mutants. (B) *NoAUREO* 3 expression in the Aureo 3-RNAi lines in the dark or after a blue light pulse, measured by RT-qPCR. The elongation factor gene (EF) was used as control. The average and range are shown (n=2). Experimental set up is the same as described in Fig. 1A but light intensity of the light pulse was 5 μmol s^−1^ m^−2^.

### Aureochromes are necessary for the blue light mediated expression of fatty acid desaturase genes

We analyzed the expression of fatty acid desaturase genes involved in EPA production in the aureochrome mutants after pulses of blue light (Fig. 6). The blue light induction of *FAD* Δ12, Δ9, Δ6, Δ5 and ω3 expression was significantly reduced in all the mutant lines analyzed when compared to the wild type. The NoAUREO 2-KO A5 line displayed the strongest reductions and showed no blue light mediated induction of *FAD* Δ9, *FAD* Δ5 and *FAD* ω3 expression. In the dark, we did not observe significant changes in the expression of the analyzed genes in the mutants, although *FAD* expression was slightly reduced in the dark samples of NoAUREO *3* RNAi lines. These results demonstrate that aureochromes are necessary for blue light expression of these genes and that these photoreceptors might influence fatty acid metabolism in *N. oceanica CCMP1779*.

**Figure 6.**
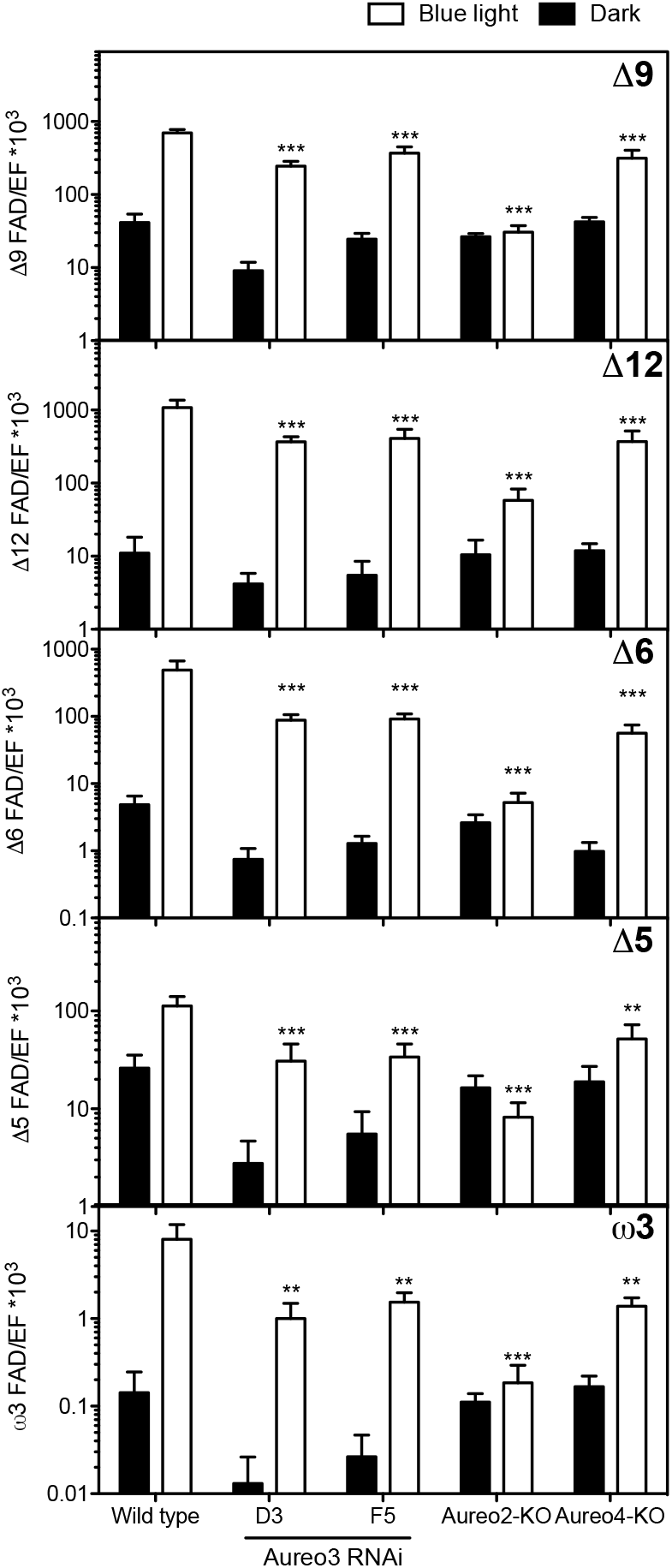
Expression of fatty acid desaturase genes in aureochrome mutants. Cells were treated as shown in Fig. 1A, the light intensity used for the light pulse was 5 μmol s^−1^ m^−2^. Gene expression was measured by RT-qPCR and the elongation factor gene (EF) was used as control. The average and standard error are shown (n=4-5). Statistical significant differences to the wild type are indicated (ANOVA and Bonferroni post hoc test; *** p< 0.001, ** p <0.01).

### Aureochromes influence growth and fatty acid profiles in *N. oceanica* CCMP1779

To investigate the role of aureochromes on fatty acid composition we measured fatty acid profiles in the aureochrome mutants (Fig. 7, Supplemental Dataset S3). We grew the cells under white light/dark cycles and then transferred them to either constant blue or red light for three days. In these experiments, growth rate under red light was lower than under blue light (p < 0.001) for all genotypes (Fig. 7A). The genotype influenced the growth rates (p < 0.001) and the aureochrome mutants displayed, in most cases, lower growth rates than the wild type line. The lines NoAUREO 2-KO #A5 and NoAUREO 4-KO #1D displayed significantly reduced growth rates with respect to the wild type (p< 0.01) under both light qualities. We did not observe a statistically significant genotype-light quality interaction affecting growth.

**Figure 7.**
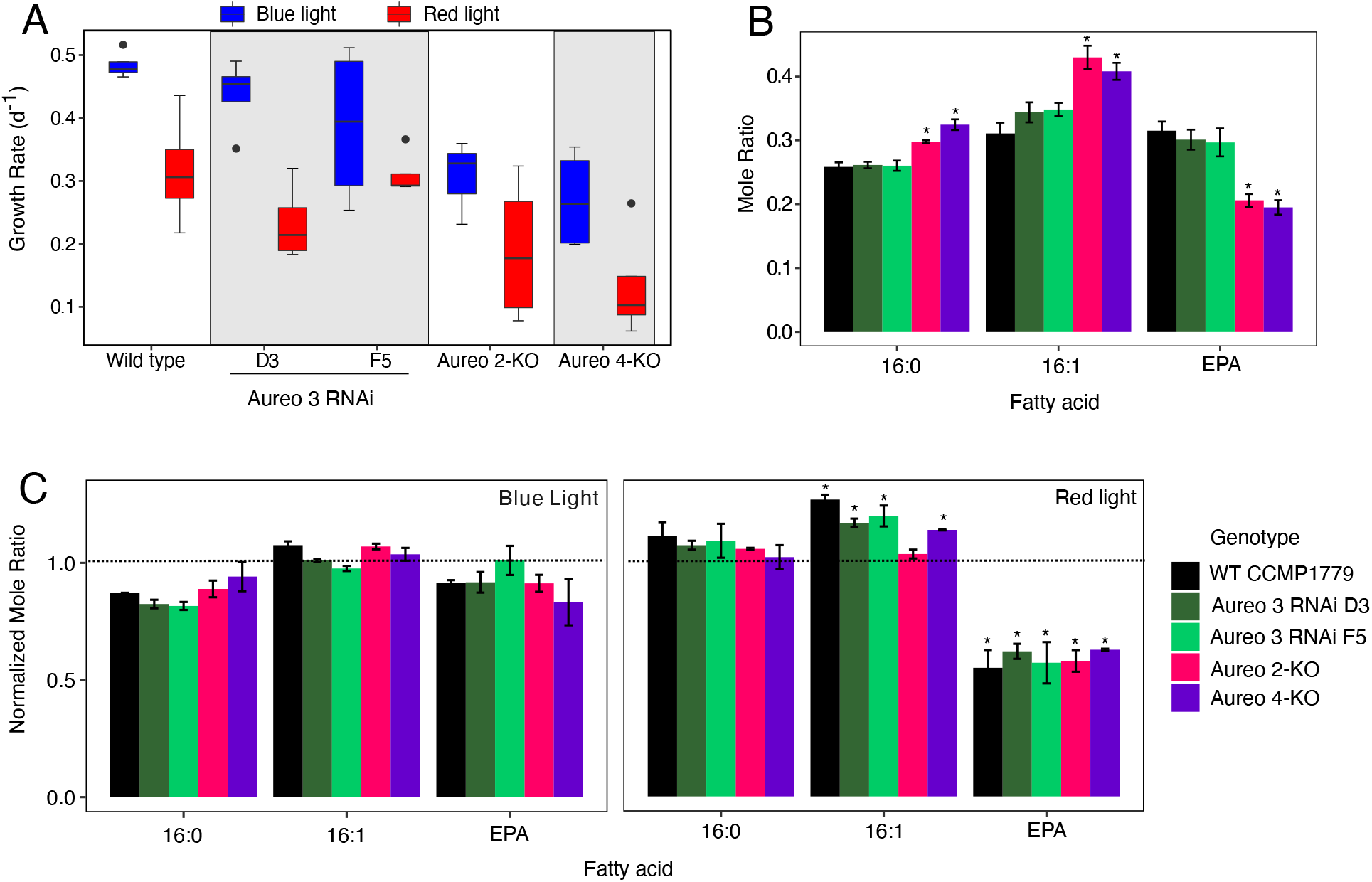
Fatty acid composition in aureochrome mutants. Cells were grown under white light/dark cycles and the transferred to either blue or red light for three days. (A) Growth rates under red or blue light, 3-4 biological replicates from 2 experiments were analyzed. (B) Mole ratio of 16:0, 16:1^Δ9^, and 20:5 (EPA) under white light/dark cycles. Data are the averages ± standard error (n=3-4). (*) Indicate significant differences to the respective wild type mole ratio (ANOVA and Tukey’s HSD, p<0.05). (C) Fatty acid mole ratio after three days under either blue or red light treatment normalized to values under white light conditions at the start of the experiment. Data are the averages ± standard error (n=3-4). (*) Indicate significant differences to values before the blue/red light treatments (ANOVA and Tukey’s HSD, p<0.05).

When grown under white light/dark cycles, the loss of *NoAUREO* 2 or *NoAUREO* 4 led to a significant decrease in EPA (20:5) and increases in the mole ratio of unsaturated fatty acid 16:0 and the monounsaturated fatty acid 16:1^Δ9^ (Fig. 7B). Although we had observed a reduction in the early blue light responsiveness of fatty acid desaturase gene expression in the NoAUREO 3 RNAi lines, these lines did not influence the percent of polyunsaturated fatty acids in these longer term experiments. However, since NoAUREO 3 RNAi lines led to an increase in 16:1 ^Δ9^ content, NoAUREO 3 might still exert some influence on the flux towards the production of polyunsaturated fatty acids.

Wild type *N. oceanica* CCMP1779 cells did not display significant changes in fatty acid profiles after being transferred from white to blue light conditions; however, transfer to red light led to a significant reduction in the EPA mole ratio (Fig. 7C) confirming our previous results (Fig. 3). We observed similar relative changes in all aureochrome mutants. Blue light did not change the fatty acid composition relative to white light, but red light led to a reduction in EPA mole ratio in all the lines tested (Fig. 5). As observed under white light, the EPA mole ratio was lower in the NoAUREO 2- and NoAUREO 4-KO lines when compared to the wild type after both blue or red light treatments (Supplemental Dataset S3), but it was not significantly changed in the NoAUREO 3 RNAi lines. These results agree with the hypothesis that blue light is required for LC-PUFA production. In the red light treatment (in the absence of blue light), EPA content is reduced over time in all genotypes as the cells grow and divide but have limited LC-PUFA biosynthesis rates.

## DISCUSSION

### Blue and red light signaling in aquatic environments

Our transcriptome analyses demonstrate that blue and red light signals trigger different but complementary responses in *N. oceanica* CCMP1779. Strong differences in gene expression have also been observed when diatoms are switched from growth in red light to blue light (Mann et al., 2020). Moreover, our analyses indicate different mechanisms of light signaling in *N. oceanica* CCMP1779. Our experiments using short pulses of blue and red light demonstrate that even in the absence of canonical red light photoreceptors, *N. oceanica* CCMP1779 was able to rapidly promote gene expression changes under red light. Most genes regulated by red light were also regulated by blue light, indicating that red light signaling is mediated by a pathway that is not wavelength specific. Our experiments using a photosynthesis inhibitor (Fig. 1D) indicate that photosynthesis might be a key driver of red light mediated changes in gene expression as has been shown to occur in diatoms (Fortunato et al., 2016). In the diatoms *P. tricornutum* and *Thalassiosira pseudonana* the far-red light sensing phytochrome DPH does not mediate red light specific changes in gene expression (Fortunato et al., 2016). Future studies using red light could help determine the role of photosynthesis signals in regulating gene expression in ochrophytes.

Sensing of blue light appears to be an ubiquitous characteristic of all photosynthetic aquatic organisms; however, photoreceptors that are able to absorb red light, such as phytochromes are less widely distributed (Duanmu et al., 2014; Duanmu et al., 2017). For example, *Nannochloropsis* species and the diatom *Thalassiosira oceanica* are some of the ochrophytes lacking phytochromes (Vieler et al., 2012; Rockwell and Lagarias, 2019). Phytochromes can act as red/far-red light photoreceptors and they appear to have an ancestral single origin among ochrophytes, and therefore, it has been proposed that their absence might have been due to a subsequent loss in some ochrophyte lineages (Fortunato et al., 2016; Rockwell and Lagarias, 2019). In land plants, red/ far red light absorbing phytochromes play key roles in detecting and responding to shading by neighbors (Roig-Villanova and Martinez-Garcia, 2016). It has been proposed that chlorophyll fluorescence and Raman scattering can generate red/far-red photons from solar green and blue light in deeper layers of the water column (Ragni and Ribera D’Alcalà, 2004; Fortunato et al., 2016). Metagenomic analyses indicate that a far-red sensing phytochrome gene is widely present in diatoms found at different depths and could mediate these signals (Fortunato et al., 2016). However, the absence of phytochromes in several ochrophyte lineages, opens the question of the adaptive role of far-red light sensing in aquatic environments.

### *N. oceanica* CCMP1779 aureochromes are able to absorb blue light and regulate blue light mediated gene expression

Low expression or loss of function mutants of any of the three aureochromes led to a reduction of FAD expression in *N. oceanica* CCMP1779 (Fig. 6). The loss of NoAUREO 2 blocked almost all early blue light responsiveness of *FAD* transcription. The blue light responsiveness of *FAD* expression was also significantly reduced in the Aureo 3 RNAi and Aureo 4-KO lines, indicating that all three aureochrome genes play partially redundant functions with respect to *FAD* regulation. Aureochromes form homo and heterodimers (Toyooka et al., 2011; Banerjee et al., 2016; Heintz and Schlichting, 2016) and one hypothesis for their partially redundant function is for *N. oceanica* CCMP1779 aureochromes to be active as heterodimers. The average expression of *NoAUREO* 4 under light/dark cycles is ~6 and ~8 times lower than that of *NoAUREO* 2 and *NoAUREO* 3 respectively (Poliner et al., 2015). The relatively low expression of *NoAUREO* 4 could explain why this gene might play less of a role in regulating the early blue light response of FAD expression. However, AUREO 2-KO and AUREO 4-KO lines displayed similar reductions in EPA content and growth rates (Fig. 7). *NoAUREO* 4 transcription is induced by light (Supplemental Fig. S5) and could be involved in a feedforward regulatory mechanism to maintain blue light regulated gene expression at longer time scales. Similarly, a recent study in the diatom *P. tricornutum* has shown that blue light also induces the expression of *PtAUREO1c*, also a class III/IV aureochrome (Supplemental Fig. S5) and that it represses the expression of *PtAUREO1b* and *PtAUREO2* (Mann et al., 2020). These changes were dependent on PtAUREO1a, indicating that stramenopile aureochromes are also involved in complex self-regulation. Transcriptome analysis of PtAureo1a mutants indicated that this aureochrome was responsible for most changes in gene expression after transfer from red to blue light, raising the question of the role of the other aureochromes in diatoms. Parallel studies on several aureochrome mutants and higher order mutants will help determine unique and redundant roles of these photoreceptors.

Our analyses of NoAUREO 3 demonstrated that the photochemistry of this protein was highly similar to previously characterized aureochromes from *V. frigida* and *P. tricornutum* (Takahashi et al., 2007; Heintz and Schlichting, 2016). The yield of NoAUREO 2 and NoAUREO 4 full length proteins when expressed in *E. coli* were significantly lower than NoAUREO 3, precluding analyses of NoAUREO 4, while allowing limited analyses of NoAUREO 2. The latter exhibited fluorescence properties similar to NoAUREO 3 in the dark-adapted state, but with higher fluorescence than NoAUREO 3 in the blue-light excited state relative to the dark state (Supplemental Fig. S7). These initial analyses indicate that NoAUREO 2 and NoAUREO 3 have the capacity to serve as canonical blue-light photoreceptors contributing to the gene regulatory responses supported by the genetic analyses of RNAi and mutant lines.

### Conservation of light regulated gene expression

Some processes are light regulated across a wide range of photosynthetic organisms, although the mechanisms of regulation are likely to be different in different groups of genes and in different organisms. In *N. oceanica* CCMP1779 both blue light and red light lead to the upregulation of translation related genes. However, functional analyses indicate that different subsets of genes are regulated by blue light signals or by the wavelength-unspecific mechanism. For example, genes involved in tRNA and amino acid activation were induced by a blue light specific mechanism and genes encoding ribosome components were induced by both blue and red light (Fig. 2). Interestingly, in the diatom *P. tricortunum*, ribosomal proteins are also induced by red light (Fortunato et al., 2016) indicating that this mechanism might be conserved among ochrophytes. Protein synthesis related genes are expressed early in the light/dark cycle in green algae and ochrophytes (Poliner et al., 2015; Zones et al., 2015; Smith et al., 2016; Carreres et al., 2019) and land plants (Michael et al., 2008) and light is necessary for translation to occur in plants (Juntawong and Bailey-Serres, 2012; Merchante et al., 2017). Translation is a high energy demanding process and is expected to be modulated by light in photosynthetic organisms.

Aureochromes and blue light have been demonstrated to regulate cell cycle progression in diatoms via modulation of the diatom specific cyclin *dsCYC2* (Huysman et al., 2013). In *N. oceanica* CCMP1779, blue light inhibited the expression of DNA replication related genes (Fig. 2) indicating that it might also work as a signal regulating the cell cycle. Although several cyclin related genes were repressed by light we identified two blue light induced cyclins (jgi|Nanoce1779_2|549456, jgi|Nanoce1779_2|667709) (Supplemental Dataset S1). The role of blue light in cell division in *Nannochloropsis* species will require future studies.

Blue light strongly and rapidly induces expression of fatty acid desaturases in *N. oceanica* CCMP1779. In our short light treatment experiments, red light also affected the expression of several of these genes, although this response was weaker than blue light (Fig. 3). Light affects the gene expression of fatty acid desaturases in other ochrophytes (Supplemental Fig. S3), and in the diatom *P. tricornutum* blue light enhances expression of these genes with respect to red light in a PtAUREO1a dependent manner indicating the same regulatory mechanism (Mann et al., 2020). Light induces FAD expression in other photosynthetic organisms. In the cyanobacterium Synechocystis PCC 6803 the induction of the transcription of fatty acid desaturase genes is dependent on photosynthesis (Kis et al., 1998). Phytochrome B is necessary for expression of fatty acid desaturases in the land plant *Arabidopsis thaliana* and shade (low red/far-red ratio) has a negative impact on fatty acid desaturation (Arico et al., 2019). Unsaturated fatty acids are mainly present in glycerolipids localized in photosynthetic membranes (Li-Beisson et al., 2019) and light signals could ensure the chloroplast growth necessary to sustain cellular growth.

## MATERIALS AND METHODS

### Strains, culture conditions and light sources used

For all experiments *N. oceanica* CCMP1779 wild type and mutant lines were grown in f/2 media and cell numbers were determined using a Beckman Coulter Z/2 using profile with a range of 1.7-4.9 μm. For the initiation of liquid cultures, cells were maintained in flasks under agitation (100 rpm), at 22 °C, under white fluorescence bulbs (100 or 200 μmol s^−1^ m^−2^) in a Percival incubator (Model GSAL-22L). Light intensity was measured using a LI-COR LI-250A Light Meter. For the light treatments, light was provided by blue or red LED lamps whose spectrum was already described (Takeuchi et al., 2014) and intensity adjusted using neutral density filters, a Heliospectra RX30 lamp using 450 nm (for blue), or white fluorescence bulbs.

### Transcriptome analyses

For each biological replicate we used 50 ml cultures grown in a 250 ml flask with a cell density of ~36-41 10^6^ cells ml^−1^ at the time of harvest. Cells were grown under 12 h light:12 h dark cycles at 22°C under agitation (100 rpm). Light intensity during light periods was 30 μmol s^−1^ m^−2^. Before harvesting, cells were transferred to darkness at ZT 12 and kept in the dark for 36 h. Cells were given a 5 min blue or red light pulse (20 μmol s^−1^ m^−2^) or kept in the dark. The spectrum of the lights was previously described (Takeuchi et al., 2014) and the intensity was adjusted using neutral density filters. All cells were then kept in the dark for 15 min before harvesting. Cells were collected at 4,000 g for 10 min at 4°C, the cell pellet was resuspended in 1.5 ml f/2 media and transferred to 2 ml tubes. The cells were centrifuged again at 18,000 g for 10 min at 4°C and flash frozen in liquid nitrogen. Black or amber tubes were used to prevent light exposure during harvesting. RNA was isolated as described previously (Poliner et al., 2015). RNA quality was checked with the Bioanalyzer (Agilent). All samples had a RIN of 7.5-8.30. Samples were sequenced at the MSU-Research Technology Support Facility using an Illumina HiSeq 2500 and a single-end 50 nucleotide run. Six samples were sequenced in each lane using a custom bar-coding. The average number of RNA-seq reads per sample was 32,000,427 and they ranged between 28,560,998 and 38,158,046.

### Analysis of differential gene expression

Read quality control and trimming was carried out using FastQC. We analyzed differential gene expression using the Tuxedo protocol (Trapnell et al., 2012). Reads were mapped to the *N. oceanica* CCMP1779 v2.0 genome annotation (JGI) using TopHat. The transcriptome was then assembled using Cufflinks. Cuffnorm was used to calculate FPKM (Fragments Per Kilobase Million) values. We determined differentially expressed genes using Cuffdiff. We defined differentially expressed genes, genes with a q-value <0.001 and either a two-fold increase to a TPM (Transcripts Per Kilobase Million) > 0.1 or a 50% decrease from a TPM > 0.1. For easier comparison with experiments using *N. oceanica* CCMP1779 V1.0 annotation we linked both annotations using BLASTn (Supplemental Dataset S4). Hierarchical cluster analysis was performed using Euclidian distances. GO term enrichment analysis was carried out using the topGO package in R. A Fisher’s exact test was used to test for significant enrichment, we defined enriched terms as having an FDR adjusted p-value < 0.05. KOG functional annotation enrichment analysis was carried out using a one-tailed Fisher’s exact test using the KOGMWU package (Dixon et al., 2015), p<0.05.

### Promoter motif analysis

Enrichment of motifs in the promoter regions of blue light induced genes was conducted using HOMER (Heinz et al., 2010) and sequences −1000 upstream from the annotated transcriptional start site of *Nannochloropsis oceanica* CCMP1779 v2.0 annotation. We selected genes that were only induced by blue light by at least four-fold (202). As background we used the respective upstream sequences off all the genes expressed (TPM > 0.1) in at least one condition in our transcriptome data. We used the command findMotifs.pl and the parameters-S 10-p 10-len 8,10.

### Expression analysis by RT-qPCR

For each biological replicate we used a 25 ml culture grown in a 50 ml flask (~15-23 10^6^ cells ml^−1^). Cells were grown under 12 h light:12 h dark cycles at 22°C under agitation (100 rpm). Light intensity during light periods was 200 μmol s^−1^ m^−2^. Before harvesting cells were transferred to darkness at ZT 12 and kept in the dark for 36 h. Cells were given a 5 min light pulse at the quality and intensity indicated in the respective figure legend or kept in the dark. All cells were then kept in the dark for 15 min before harvesting. For the DCMU treatments, cells were treated with either 5 μl of a 100 mM DCMU in ethanol for a final concentration of 20 μM, or with 5 μl ethanol as control 10 min before the light pulse. Cells were collected at 4,000 g for 5-10 min at 17°C, the cell pellet was resuspended in 1.5 ml f/2 media and transferred to 2 ml tubes. The cells were centrifuged again at 5,000 g for 4 min at 4°C and flash frozen in liquid nitrogen. Black or amber tubes were used to prevent light exposure during harvesting. RNA was isolated as described previously (Poliner et al., 2015). DNA was removed from RNA extracts using Turbo DNase (Thermo Fisher Scientific) according to the manufacturer recommendations. cDNA synthesis and RT-qPCR was performed as described previously (Poliner et al., 2015). The genes ACTR (jgi|Nanoce1779_2|596289) or EF (jgi|Nanoce1779_2|544329) were used as control. Primers used are on Table S1.

### Oxygen measurements

For each biological replicate we used a 25 ml culture grown in a 50 ml flask (~25-29 10^6^ cells ml^−1^). Cells were grown under 12 h light:12 h dark cycles at 22°C under agitation (100 rpm). Light intensity during light periods was 200 μmol s^−1^ m^−2^. Oxygen was measured using a Hansatech oxygraphy system (Hansatech, Norfolk, UK). Cells were illuminated using a F&V HD R-300 white ring LED light (F&V lighting, Mundelein, IL, USA) and the intensity at cell level was 34 μmol s^−1^ m^−2^. Cells (2ml) were transferred to the chamber and treated with either 0.4 μl of a 100 mM DCMU/ethanol solution (final concentration of 20 μM), or with 0.4 μl ethanol as vehicle control 10 min before the start of measurements.

### Cloning of Aureochrome coding regions

Based on *N. oceanica* CCMP1779 v1.0 and v2.0 annotations and aligned sequencing reads we cloned and sequenced the CDS of three aureochrome genes (Supplemental Dataset S2). The coding region for *NoAUREO* 4 (protein ID v1.0: NannoCCMP1779|5385) corresponds best with the v1.0 annotation, which also matches best the aligned transcriptome reads. The current corresponding v2.0 annotation protein ID is jgi|Nanoce1779_2|622551. The coding region we cloned for *NoAUREO* 3 lacks the annotated N-terminal extension in both annotations (protein ID for v2.0: jgi|Nanoce1779_2|582424; for v1.0: NannoCCMP1779|10447), but the region we were able to clone matches well with the RNA-seq reads for *N. oceanica* CCMP1779. The cloned coding region for *NoAUREO* 2 corresponds to the v2.0 annotation (protein ID jgi|Nanoce1779_2|550823). For references, in the corresponding protein ID in the v1.0 annotation is NannoCCMP1779|10112. We amplified the corresponding cDNAs using the primers on Table S1 and Q5^®^ High-Fidelity DNA Polymerase (NEB). The PCR fragments were cloned into pENTR-D Topo vector and sequenced.

### Cloning of NoAUREO 3 RNAi constructs

For the generation of the aureochrome 3 RNAi lines we used the plasmid pNOC-642 (Poliner et al., 2020), that expresses the RNAi construct under the control of the *N. oceanica* CCMP1779 bidirectional Ribi promoter (Poliner et al., 2018) coexpressed with the Nourseothricin resistance gene. We amplified the sense *NoAUREO* 3 fragment from cDNA using the primers A3-sense-F and A3-sense-R (Table S1) and the antisense strand using A3-antisense-F and A3-antisense-R, using Q5^®^ High-Fidelity DNA Polymerase (NEB). The vector pNOC-642 linearized using SacI (NEB) was assembled with the sense and antisense fragments using NEBuilder^®^ HiFi DNA Assembly (NEB) according to the manufacturer instructions, so that the antisense construct was cloned between the Ribi promoter and the LDSP terminator. The plasmid was checked by sequencing and used to transform *N. oceanica* CCMP1779 cells as described in (Poliner et al., 2018). Transformants were plated on f/2 plates supplemented with nourseothricin (200 μg ml^−1^) under grown under constant light for three weeks.

### Generation of knock out mutants using CRISPR-Cas9

For the creation of aureochrome targeting CRISPR constructs guide sequences to target the aureochrome genes were generated according to the strategy described previously (Poliner et al., 2018). To avoid frameshift mutations, we used tandem sgRNAs targeting both ends to the aureochrome genes.

The pNOC-ARS-CRISPR-compact (Addgene: 99369) (Poliner et al., 2018) vector was modified to tie resistance marker gene expression directly to Cas9-Nlux expression by the P2A peptide linker to generate the plasmid pNOC-ARS-CRISPR-P2A-BlastR (Addgene: 98147). The P2A-BlastR fragment was amplified from pNOC-ARS-destiny-P2A-BlastR-GFP (Poliner et al., 2020) using the primers SV40 P2A nhei F+/ BlastR mfei R- (Table S1). The construct pNOC-ARS-CRISPR-compact and the PCR product was digested with NheI/MfeI (NEB) and ligated to generate the vector pNOC-ARS-CRISPR-P2A-BlastR (Supplemental Dataset S2).

Tandem sgRNAs were assembled in a vector through a two-step cloning strategy. The sgRNAs targeting the 3’ end of *AUREO* 2 and *AUREO* 4 genes were created by site directed mutagenesis using the primers sgAureo#-end F+/R- and pNOC-ARS-CRISPR-HA as template (Table S1). After ligation, sanger sequencing of these sgRNA subcloning vectors verified the correct sequences. The sgRNA subcloning vectors were then used as templates in PCR amplification with the primers sgHH BstBI and sgHH HDV ClaI KpnI R-. This generated a PCR amplicon with the 5’ ClaI site changed into a compatible BstBI site and also added 3’ ClaI and KpnI restriction sites. This PCR fragment was digested with BstBI/KpnI and the vector pNOC-ARS-CRISPR-P2A-BlastR was digested with ClaI/KpnI. Ligation resulted in the destruction of the 5’ BstBI restriction site and unique ClaI and KpnI sites 3’ of the sgRNA. The sgRNAs targeting the 5’ ends of the genes were created by site directed mutagenesis using the primers sghh Aureo2-1 F+/R- and sghh Aureo4-3 F+/R-primers with the vectors pNOC-ARS-CRISPR-HA (*AUREO* 2) or pNOC-CRISPR-PSPX-U6 (*AUREO* 4) as template (Table S1). The sgRNA sequences were verified by Sanger sequencing of the sgRNA subcloning vectors. The 5’ targeting sgRNAs were transferred to the pNOC-ARS-CRISPR-P2A-BlastR plasmids, already containing the respective sgRNAs targeting the 3’ end of the genes, by ClaI/KpnI digestion and ligation. In order to maintain constructs as episomes, the circular plasmids were introduced into *N. oceanica* CCMP1779 cells as described previously (Poliner et al., 2018).

Resistant colonies were grown in 96 deep-well plates in f/2 supplemented with blasticidin (50 μg ml^−1^). Transformants were screened for luminescence to indicate Cas9-Nlux presence (Poliner et al., 2018). Following Cas9-Nlux detection selected transformants were screened with PCR. Primers (pro seq F+/term seq R-) were placed approximately 500 base pairs from the target sites and an extension time sufficient for the full length genomic sequence was used. Deletions were apparent from decrease in the size of the amplicon. The exact mutations were determined by Sanger sequencing using internal primers (target seq F+) (Table S1).

### Protein expression and purification

For expression in *E. coli* NoAureo2 was cloned using pENTR NoAUREO 2 as a template with the following primers (Table S1): Aureo2(BamH1)fwd and Aureo2(EcoR1)rev. Aureo3 was cloned using pENTRaureo3 as a template with the following primers (Table S1): Aureo3(BamH1)fwd and Aureo3(EcoR1)rev. Aureo4 was cloned using pENTR NoAUREO4 as a template with the following primers (Table S1): Aureo4(BamH1)fwd and Aureo4(EcoR1)rev. *NoAUREO* 2, *NoAUREO* 3, and *NoAUREO* 4 were amplified using 30 cycles of 98 °C for 10 sec, 55 °C for 5 sec, and 72 °C for 5 sec. The PCR-products (*NoAUREO* 2, 1140 bp; *NoAUREO* 3, 720 bp; and *NoAUREO* 4, 1470 bp) were purified and restricted with BamH1 and EcoR1 prior to subcloning into pGEX-6P-1 (GE Healthcare, Uppsala, Sweden) digested with the same enzymes to produce plasmids to produced N-terminally GST-tagged versions of NoAUREO 2, NoAUREO 3, and NoAUREO 4 proteins.

Protein expression and purification: pGEX6P1_aureo2, pGEX6P1_aureo3, and pGEX6P1_aureo4 in BL21 cells were grown in autoinduction medium containing ampicillin (100 μg/mL) at 17° C for 72 h. Cell pellets were collected for 250 ml of induced culture at 8400 *g* at 4° C for 15 min. Pellet cells were resuspended in 10 mL of 20 mM Tris, 400 mM NaCl at pH 7.4 with the addition of 0.2 mg/mL (w/v) lysozyme, 2 μL or 500 U Benzonase^®^ Endonuclease (Sigma-Aldrich), 1.25 mM EDTA, and 1 mM DTT. Resuspended cells passed through a French Press at working pressure of ~20K psi twice. The lysate was spun at 17,000 g at 4 °C for 30 min to collect the soluble fraction.

A 400 μL aliquot of Pierce^™^ Glutathione Agarose (Pierce) was used for column-based purification of the recombinant aureochrome proteins. A packed column was equilibrated at 5 °C with 10 column volumes of a Tris-based buffer containing 20 mM Tris, 400 mM NaCl, 1 mM DTT at pH of 7.4. A 5 mL aliquot of soluble protein extract was diluted with 10 mL of the Tris buffer used to equilibrate the column and the total 15 mL sample added to the column. After the sample flowed through the column based on gravity flow, the column was washed with 10 ml of the Tris-based buffer. The GST tag was cleaved from purified protein by elution of the tag-free protein from the column by overnight incubation with PreScission Protease (GE Healthcare) or alternatively by elution with 15 mM glutathione-containing purification buffer, followed by overnight dialysis with PreScission Protease and subsequent removal of GST and the protease by a second affinity purification. Cleaved protein was dialyzed against 20 mM Tris, 200 mM NaCl, 1 mM DTT at pH of 7.4.

### Spectroscopic analyses

Absorbance and fluorescence spectra were measured at room temperature using dialyzed protein. The protein concentration was ~20 μM in 20 mM Tris, 200 mM NaCl, 1 mM DTT at pH of 7.4. Protein samples were illuminated in a quartz cuvette with a path length of 1 cm with a blue light (LED, 450 nm) for 1 min where indicated. Dark recovery of excited samples was conducted by transferring blue light-excited protein to dark and conducting an absorbance spectral scan from 300 nm to 550 nm using an Agilent HP8453 UV/visible spectrophotometer (Agilent Technologies) or detecting recovery of fluorescence from 470 to 595 nm using a PTI QuantaMaster spectrofluorimeter (HORIBA Instruments, Edison, NJ).

### Statistical analysis

Two-way ANOVA tests followed by Bonferroni or Tukey’s HSD *post hoc* tests were implemented in either Prism GraphPad or R.

### Mutant growth and fatty acid analyses

Mutant and wild type cells were grown under 12 h light/12 h dark in white light (30 μmol s^−1^ m^−2^) for 3-4 days. At 4-5 h after dawn they were diluted to 10-17 10^6^ cells ml^−1^. Two 2 ml aliquots were collected using GF/C glass microfiber filters (20 mm; Whatman, GE Healthcare Life Sciences), transferred to glass tubes, frozen in liquid nitrogen and kept at −80 °C. Cells were then transferred to either constant blue (34 μmol s^−1^ m^−2^) or red light (35 μmol s^−1^ m^−2^) for four days. Every 24 h, cells were counted and aliquots were taken for fatty acid analyses. Fatty acid methyl ester (FAME) extraction and analysis were carried out as described previously (Liu et al., 2013).

### Accession Numbers

*Nannochloropsis oceanica* CCMP1779 genome data and transcript or protein IDs are from *Nannochloropsis oceanica* CCMP1779 v2.0 if not indicated otherwise (JGI, https://mycocosm.jgi.doe.gov/Nanoce1779_2/Nanoce1779_2.home.html). In the main text, gene IDs from this annotation are indicated as jgi|Nanoce1779_2|XXXXX. As needed the v1.0 annotation was also used, and IDs of from this annotation are indicated as NannoCCMP1779|XXXXX. For comparison with experiments using *N. oceanica* CCMP1779 v1.0 annotation (Vieler et al., 2012)(https://mycocosm.jgi.doe.gov/Nanoce1779/Nanoce1779.home.html) we linked both annotations using BLASTp and BLASTn (Supplemental Dataset S4). Protein IDs for the fatty acid desaturases are: *Δ*9 FAD (protein ID: jgi|Nanoce1779_2|593563), *Δ*6 FAD (protein ID: jgi|Nanoce1779_2|622551), *Δ*12 FAD (jgi|Nanoce1779_2|591534), *Δ*5 FAD (jgi|Nanoce1779_2|593992) and *ω3* FAD (jgi|Nanoce1779_2|639463). Raw read data have been deposited in NCBI’s Gene Expression Omnibus (Edgar et al., 2002) and are accessible through GEO Series accession number GSE164714. For species other than *Nannochloropsis*, IDs are from GenBank, JGI or http://nandesyn.single-cell.cn, as indicated.

## Author contributions

E.M.F and E.P conceived the project. E.P., L.N., A.W.U.B., R.C., S.C.G-M and E.M.F performed the experiments. A.W.U.B. and B.L.M. designed the photochemistry experiments. Y.U.K, E.P. and E.M.F. analyzed the transcriptome data. B.L.M., B.J.J. and E.M.F. supervised the experiments. E.M.F. and wrote the article with contributions of all the authors. E.M.F. agrees to serve as the author responsible for contact and ensures communication.

## Acknowledgments

This work was funded by grant NSF IOS-1354721 to EMF and funds from Michigan State University; general infrastructure support was provided by the U.S. Department of Energy [DE-FG02-91ER20021] to BLM. SG-M was supported by the PlantGenomics@MSU program [NSF DBI-1358474]. We thank Ron Cook and Christoph Benning (Michigan State University) for help with the fatty acid analyses, and Han Bao and Berkley Walker (Michigan State University) for help with the oxygen measurements.

---

**Figure S1.** Hierarchical cluster analysis of the log transformed TPM values of differentially expressed genes in each sample.

**Figure S2. Oxygen production after DCMU treatment. Cells were grown under light/dark cycles.** During the light period cell aliquots were taken, treated with either ethanol (vehicle control) or DCMU for 10 minutes, and the rate of oxygen concentration change was measured at 34 μmol s^−1^ m^−2^ white light using an oxygen electrode. Values are the average and standard error (n=3).

**Figure S3.** Enrichment analyses of EuKaryotic Orthologous Groups (KOG). Only categories with significantly enriched functional terms are shown.

**Figure S4. Expression of fatty acid desaturase genes in *Phaeodactylum tricornutum***. Data are from (Chauton et al., 2013; Smith et al., 2016). Proposed biochemical functions are from (Dolch and Marechal, 2015) or from genome annotation. White bars indicate light period, black and grey bars indicate dark period in the Smith et al and Chauton et al experiments respectively. (*) Indicates red light induction of at least two-fold (Fortunato et al., 2016).

**Figure S5. Aureochrome proteins of N. oceanica CCMP1779.** (A) Schematic representation of the domain structure of Aureo 2, 3 and 4. Domain positions were designated based on SMART 10. The PAS and PAC domains were designated as LOV domain here. The average theoretical molecular weights as computed by Expasy Compute pI /Mw Tool are 40,828.65 (NoAUREO 2), 25,949.18 (NoAUREO 3) and 52,370.26 (NoAUREO 4). (B) Phylogenetic analysis of Aureochrome bZIP & LOV domains. Analysis were done using MEGA X (Stecher et al., 2020). Proteins were aligned using Muscle and trimmed to include only the bZIP and LOV domains. The evolutionary history was inferred using the Neighbor-Joining method (Saitou and Nei, 1987). The optimal tree with the sum of branch length = 5.65909394 is shown. The percentage of replicate trees in which the associated taxa clustered together in the bootstrap test (1000 replicates) are shown next to the branches (Jones et al., 1992). The tree is drawn to scale, with branch lengths in the same units as those of the evolutionary distances used to infer the phylogenetic tree. The evolutionary distances were computed using the JTT matrix-based method (Kumar et al., 2018) and are in the units of the number of amino acid substitutions per site. Cm, *Chattonella marina*; Es, *Ectocarpus siliculosus*; Ng, *Nanochloropsis gaditana* CCMP526; Ns, *N. salina* CCMP1776; No, *N. oceanica* CCMP1779; Od, *Ochromonas Danica*; Pt, *Phaeodactylum tricornutum*; To, *Thalassiosira oceanica*; Vf, *Vaucheria frigida*. IDs are from GenBank with the exception of *P. tricornutum* Phatr2 (JGI) and *N. salina* CCMP1776 (http://nandesyn.single-cell.cn,v2.0). The cloned coding regions for *N. oceanica* CCMP1779 (Dataset S2) were used for this analysis. Aureochrome ‘types’ are based on published classification (Schellenberger Costa et al., 2013).

**Figure S6. Light quality influences aureochrome expression in *N. oceanica* CCMP1779**. Expression of fatty acid desaturase genes after pulses of red, blue or white light. Cells were grown under light/dark cycles, transferred to 36 h of constant darkness at ZT12 and given a 5 min light pulse. Cells were collected 15 min after the pulse ended. Cells kept in the dark were used as controls. (A) Data from the RNA-seq experiment shown as log2 transformed fold change in expression with respect to the dark treated samples. Light intensity during the light pulse was 20 μmol s^−1^ m^−2^ (B) A separate experiment measured gene expression changes using RT-qPCR, results are shown as fold change with respect to the dark control. Light intensity during the light pulse was 20 μmol s^−1^ m^−2^. The actin-related gene (*ACTR*) was used as control. The average and standard error are shown (n=3).

**Figure S7. Recombinant NoAUREO expression.** (A) E. coli cells (BL21) expressing NoAUREO 3 proteins. Cell pellets were collected from cultures grown in LB, SB, or autoinduction (AI) medium containing ampicillin (100 μg/mL) at 15° C for 72 h. (B) E. coli cells (BL21) expressing NoAUREO 2 or NoAUREO 4 proteins. Cell pellets were collected from cultures grown in autoinduction (AI) medium containing ampicillin (100 μg/mL) at 15° C for 72 h.

**Figure S8. Fluorescence spectrum of recombinant NoAUREO 2**. Fluorescence spectrum of recombinant protein in the dark state or after 1 min of blue light treatment.

**Table S1.** Primers used in this study.

**Dataset S1.** Expression level and functional annotation of *N. oceanica* CCMP1779 genes after a pulse of either blue or red light.

**Dataset S2.** (A) HOMER output of motif enrichment in blue light induced genes. Regions −1000 bp from the transcriptional start site of genes induced at least four-fold in only blue light treatment were compared to the upstream regions of all genes expressed in at least one condition in the experiment. (B) Cloned coding regions of *N. oceanica* CCMP1779 aureochromes and their respective protein sequences. cDNA was used to amplify these sequences using primers on Table S1. (C) Description of the Aureochrome mutants and RNAi lines in the context of their annotation. For *NoAUREO* 2 and *NoAUREO* 4, the description includes the sequenced regions flanking the deletions and the sgRNA target sequence used. For *NoAUREO* 3 RNAi, the RNAi target is indicated. (D) Vectors used during the generation of CRISPR-Cas9 episomes.

**Dataset S3.** Fatty acid mole ratios in *N. oceanica* CCMP1779 Aureochrome mutants. Cells were grown under white light/dark cycles and transferred on day 0 to either constant blue (B) or red (L) light for three days. Data represents the average mole ratio and standard error (SE) of 3-4 biological replicates from 2 independent experiments.

**Dataset S4.** Associations between *N. oceanica* CCMP1779 v0.1 and v0.2 annotations. Protein and transcript sequences between both annotations were compared using BLASTp and BLASTn respectively.

## Parsed Citations

**Adarme-Vega TC, Thomas-Hall SR, Schenk PM (2014) Towards sustainable sources for omega-3 fatty acids production. Curr Opin Biotechnol 26: 14-18**

Google Scholar: <u>Author Only</u> <u>Title Only</u> <u>Author and Title</u>

**Alboresi A, Perin G, Vitulo N, Diretto G, Block M, Jouhet J, Meneghesso A, Valle G, Giuliano G, Marechal E, Morosinotto T (2016) Light Remodels Lipid Biosynthesis in Nannochloropsis gaditana by Modulating Carbon Partitioning between Organelles. Plant Physiol 171: 2468-2482**

Google Scholar: <u>Author Only</u> <u>Title Only</u> <u>Author and Title</u>

**Arico D, Legris M, Castro L, Garcia CF, Laino A, Casal JJ, Mazzella MA (2019) Neighbour signals perceived by phytochrome B increase thermotolerance in Arabidopsis. Plant Cell Environ 42: 2554-2566**

Google Scholar: <u>Author Only</u> <u>Title Only</u> <u>Author and Title</u>

**Banerjee A, Herman E, Serif M, Maestre-Reyna M, Hepp S, Pokorny R, Kroth PG, Essen LO, Kottke T (2016) Allosteric communication between DNA-binding and light-responsive domains of diatom class I aureochromes. Nucleic Acids Res 44: 5957-5970**

Google Scholar: <u>Author Only</u> <u>Title Only</u> <u>Author and Title</u>

**Carreres BM, León-Saiki GM, Schaap PJ, Remmers IM, van der Veen D, Martins dos Santos VAP, Wijffels RH, Martens DE, Suarez-Diez M (2019) The diurnal transcriptional landscape of the microalga Tetradesmus obliquus. Algal Research 40: 101477**

Google Scholar: <u>Author Only</u> <u>Title Only</u> <u>Author and Title</u>

**Chauton MS, Winge P, Brembu T, Vadstein O, Bones AM (2013) Gene Regulation of Carbon Fixation, Storage, and Utilization in the Diatom Phaeodactylum tricornutum Acclimated to Light/Dark Cycles. Plant Physiology 161: 1034-1048**

Google Scholar: <u>Author Only</u> <u>Title Only</u> <u>Author and Title</u>

**Chen C-Y, Chen Y-C, Huang H-C, Ho S-H, Chang J-S (2015) Enhancing the production of eicosapentaenoic acid (EPA) from Nannochloropsis oceanica CY2 using innovative photobioreactors with optimal light source arrangements. Bioresource technology 191: 407-413**

Google Scholar: <u>Author Only</u> <u>Title Only</u> <u>Author and Title</u>

**Chen C-Y, Chen Y-C, Huang H-C, Huang C-C, Lee W-L, Chang J-S (2013) Engineering strategies for enhancing the production of eicosapentaenoic acid (EPA) from an isolated microalga Nannochloropsis oceanica CY2. Bioresource Technology 147: 160-167**

Google Scholar: <u>Author Only</u> <u>Title Only</u> <u>Author and Title</u>

**Coesel S, Mangogna M, Ishikawa T, Heijde M, Rogato A, Finazzi G, Todo T, Bowler C, Falciatore A (2009) Diatom PtCPF1 is a new cryptochrome/photolyase family member with DNA repair and transcription regulation activity. EMBO Rep 10: 655-661**

Google Scholar: <u>Author Only</u> <u>Title Only</u> <u>Author and Title</u>

**Das P, Lei W, Aziz SS, Obbard JP (2011) Enhanced algae growth in both phototrophic and mixotrophic culture under blue light.**

**Bioresour Technol 102: 3883-3887**

Google Scholar: <u>Author Only</u> <u>Title Only</u> <u>Author and Title</u>

**Dixon GB, Davies SW, Aglyamova GA, Meyer E, Bay LK, Matz MV (2015) Genomic determinants of coral heat tolerance across latitudes. Science 348: 1460-1462**

Google Scholar: <u>Author Only</u> <u>Title Only</u> <u>Author and Title</u>

**Dolch LJ, Marechal E (2015) Inventory of fatty acid desaturases in the pennate diatom Phaeodactylum tricornutum. Mar Drugs 13: 1317-1339**

Google Scholar: <u>Author Only</u> <u>Title Only</u> <u>Author and Title</u>

**Duanmu D, Bachy C, Sudek S, Wong CH, Jimenez V, Rockwell NC, Martin SS, Ngan CY, Reistetter EN, van Baren MJ, Price DC, Wei CL, Reyes-Prieto A, Lagarias JC, Worden AZ (2014) Marine algae and land plants share conserved phytochrome signaling systems. Proc Natl Acad Sci U S A 111: 15827-15832**

Google Scholar: <u>Author Only</u> <u>Title Only</u> <u>Author and Title</u>

**Duanmu D, Rockwell NC, Lagarias JC (2017) Algal light sensing and photoacclimation in aquatic environments. Plant Cell Environ 40: 2558-2570**

Google Scholar: <u>Author Only</u> <u>Title Only</u> <u>Author and Title</u>

**Edgar R, Domrachev M, Lash AE (2002) Gene Expression Omnibus: NCBI gene expression and hybridization array data repository. Nucleic Acids Res 30: 207-210**

Google Scholar: <u>Author Only</u> <u>Title Only</u> <u>Author and Title</u>

**Essen LO, Franz S, Banerjee A (2017) Structural and evolutionary aspects of algal blue light receptors of the cryptochrome and aureochrome type. J Plant Physiol 217: 27-37**

Google Scholar: <u>Author Only</u> <u>Title Only</u> <u>Author and Title</u>

**Fawley KP, Fawley MW (2007) Observations on the diversity and ecology of freshwater Nannochloropsis (Eustigmatophyceae), with descriptions of new taxa. Protist 158: 325-336**

Google Scholar: <u>Author Only</u> <u>Title Only</u> <u>Author and Title</u>

**Fortunato AE, Jaubert M, Enomoto G, Bouly JP, Raniello R, Thaler M, Malviya S, Bernardes JS, Rappaport F, Gentili B, Huysman MJ, Carbone A, Bowler C, d’Alcala MR, Ikeuchi M, Falciatore A (2016) Diatom Phytochromes Reveal the Existence of Far-Red-Light-Based Sensing in the Ocean. Plant Cell 28: 616-628**

Google Scholar: <u>Author Only</u> <u>Title Only</u> <u>Author and Title</u>

**Heintz U, Schlichting I (2016) Blue light-induced LOV domain dimerization enhances the affinity of Aureochrome 1a for its target DNA sequence. Elife 5: e11860**

Google Scholar: <u>Author Only</u> <u>Title Only</u> <u>Author and Title</u>

**Heinz S, Benner C, Spann N, Bertolino E, Lin YC, Laslo P, Cheng JX, Murre C, Singh H, Glass CK (2010) Simple combinations of lineage-determining transcription factors prime cis-regulatory elements required for macrophage and B cell identities. Mol Cell 38: 576-589**

Google Scholar: <u>Author Only</u> <u>Title Only</u> <u>Author and Title</u>

**Hisatomi O, Takeuchi K, Zikihara K, Ookubo Y, Nakatani Y, Takahashi F, Tokutomi S, Kataoka H (2013) Blue light-induced conformational changes in a light-regulated transcription factor, aureochrome-1. Plant Cell Physiol 54: 93-106**

Google Scholar: <u>Author Only</u> <u>Title Only</u> <u>Author and Title</u>

**Huysman MJ, Fortunato AE, Matthijs M, Costa BS, Vanderhaeghen R, Van den Daele H, Sachse M, Inze D, Bowler C, Kroth PG, Wilhelm C, Falciatore A, Vyverman W, De Veylder L (2013) AUREOCHROME1a-mediated induction of the diatom-specific cyclin dsCYC2 controls the onset of cell division in diatoms (Phaeodactylum tricornutum). Plant Cell 25: 215-228**

Google Scholar: <u>Author Only</u> <u>Title Only</u> <u>Author and Title</u>

**Jones DT, Taylor WR, Thornton JM (1992) The rapid generation of mutation data matrices from protein sequences. Comput Appl Biosci 8: 275-282**

Google Scholar: <u>Author Only</u> <u>Title Only</u> <u>Author and Title</u>

**Juntawong P, Bailey-Serres J (2012) Dynamic Light Regulation of Translation Status in Arabidopsis thaliana. Front Plant Sci 3: 66**

Google Scholar: <u>Author Only</u> <u>Title Only</u> <u>Author and Title</u>

**Kis M, Zsiros O, Farkas T, Wada H, Nagy F, Gombos Z (1998) Light-induced expression of fatty acid desaturase genes. Proc Natl Acad Sci U S A 95: 4209-4214**

Google Scholar: <u>Author Only</u> <u>Title Only</u> <u>Author and Title</u>

**Kroth PG, Wilhelm C, Kottke T (2017) An update on aureochromes: Phylogeny - mechanism - function. J Plant Physiol 217: 20-26**

Google Scholar: <u>Author Only</u> <u>Title Only</u> <u>Author and Title</u>

**Kumar S, Stecher G, Li M, Knyaz C, Tamura K (2018) MEGAX: Molecular Evolutionary Genetics Analysis across Computing Platforms. Mol Biol Evol 35: 1547-1549**

Google Scholar: <u>Author Only</u> <u>Title Only</u> <u>Author and Title</u>

**Li-Beisson Y, Thelen JJ, Fedosejevs E, Harwood JL (2019) The lipid biochemistry of eukaryotic algae. Prog Lipid Res 74: 31-68**

Google Scholar: <u>Author Only</u> <u>Title Only</u> <u>Author and Title</u>

**Liu B, Vieler A, Li C, Daniel Jones A, Benning C (2013) Triacylglycerol profiling of microalgae Chlamydomonas reinhardtii and Nannochloropsis oceanica. Bioresour Technol 146: 310-316**

Google Scholar: <u>Author Only</u> <u>Title Only</u> <u>Author and Title</u>

**Locke JC, Southern MM, Kozma-Bognar L, Hibberd V, Brown PE, Turner MS, Millar AJ (2005) Extension of a genetic network model by iterative experimentation and mathematical analysis. Mol Syst Biol 1: 2005.0013**

Google Scholar: <u>Author Only</u> <u>Title Only</u> <u>Author and Title</u>

**Mann M, Serif M, Jakob T, Kroth PG, Wilhelm C (2017) PtAUREO1a and PtAUREO1b knockout mutants of the diatom Phaeodactylum tricornutum are blocked in photoacclimation to blue light. J Plant Physiol 217: 44-48**

Google Scholar: <u>Author Only</u> <u>Title Only</u> <u>Author and Title</u>

**Mann M, Serif M, Wrobel T, Eisenhut M, Madhuri S, Flachbart S, Weber APM, Lepetit B, Wilhelm C, Kroth PG (2020) The Aureochrome Photoreceptor PtAUREO1a Is a Highly Effective Blue Light Switch in Diatoms. iScience 23: 101730**

Google Scholar: <u>Author Only</u> <u>Title Only</u> <u>Author and Title</u>

**Merchante C, Stepanova AN, Alonso JM (2017) Translation regulation in plants: an interesting past, an exciting present and a promising future. Plant J 90: 628-653**

Google Scholar: <u>Author Only</u> <u>Title Only</u> <u>Author and Title</u>

**Michael TP, Mockler TC, Breton G, McEntee C, Byer A, Trout JD, Hazen SP, Shen R, Priest HD, Sullivan CM, Givan SA, Yanovsky M, Hong F, Kay SA, Chory J (2008) Network discovery pipeline elucidates conserved time-of-day-specific cis-regulatory modules. PLoS Genet 4: e14**

Google Scholar: <u>Author Only</u> <u>Title Only</u> <u>Author and Title</u>

**Poliner E, Clark E, Cummings C, Benning C, Farre EM (2020) A high-capacity gene stacking toolkit for the oleaginous microalga, Nannochloropsis oceanica CCMP1779. Algal Research 45: 101664**

Google Scholar: <u>Author Only</u> <u>Title Only</u> <u>Author and Title</u>

**Poliner E, Farre EM, Benning C (2018) Advanced genetic tools enable synthetic biology in the oleaginous microalgae Nannochloropsis sp. Plant Cell Rep 37: 1383-1399**

Google Scholar: <u>Author Only</u> <u>Title Only</u> <u>Author and Title</u>

**Poliner E, Panchy N, Newton L, Wu G, Lapinsky A, Bullard B, Zienkiewicz A, Benning C, Shiu SH, Farre EM (2015) Transcriptional coordination of physiological responses in Nannochloropsis oceanica CCMP1779 under light/dark cycles. Plant Journal 83: 1097-1113**

Google Scholar: <u>Author Only</u> <u>Title Only</u> <u>Author and Title</u>

**Poliner E, Pulman JA, Zienkiewicz K, Childs K, Benning C, Farre EM (2018) A toolkit for Nannochloropsis oceanica CCMP1779 enables gene stacking and genetic engineering of the eicosapentaenoic acid pathway for enhanced long-chain polyunsaturated fatty acid production. Plant Biotechnol J 16: 298-309**

Google Scholar: <u>Author Only</u> <u>Title Only</u> <u>Author and Title</u>

**Poliner E, Takeuchi T, Du ZY, Benning C, Farre EM (2018) Nontransgenic Marker-Free Gene Disruption by an Episomal CRISPR System in the Oleaginous Microalga, Nannochloropsis oceanica CCMP1779. ACS Synth Biol 7: 962-968**

Google Scholar: <u>Author Only</u> <u>Title Only</u> <u>Author and Title</u>

**Ragni M, Ribera D’Alcalà M (2004) Light as an information carrier underwater. Journal of Plankton Research 26: 433-443**

Google Scholar: <u>Author Only</u> <u>Title Only</u> <u>Author and Title</u>

**Rockwell NC, Lagarias JC (2019) Phytochrome evolution in 3D: deletion, duplication, and diversification. New Phytol 225: 2283–2300**

Google Scholar: <u>Author Only</u> <u>Title Only</u> <u>Author and Title</u>

**Roig-Villanova I, Martinez-Garcia JF (2016) Plant Responses to Vegetation Proximity: A Whole Life Avoiding Shade. Front Plant Sci 7: 236**

Google Scholar: <u>Author Only</u> <u>Title Only</u> <u>Author and Title</u>

**Saitou N, Nei M (1987) The neighbor-joining method: a new method for reconstructing phylogenetic trees. Mol Biol Evol 4: 406-425**

Google Scholar: <u>Author Only</u> <u>Title Only</u> <u>Author and Title</u>

**Sayanova O, Mimouni V, Ulmann L, Morant-Manceau A, Pasquet V, Schoefs B, Napier JA (2017) Modulation of lipid biosynthesis by stress in diatoms. Philos Trans R Soc Lond B Biol Sci 372: 20160407**

Google Scholar: <u>Author Only</u> <u>Title Only</u> <u>Author and Title</u>

**Schellenberger Costa B, Sachse M, Jungandreas A, Bartulos CR, Gruber A, Jakob T, Kroth PG, Wilhelm C (2013) Aureochrome 1a is involved in the photoacclimation of the diatom Phaeodactylum tricornutum. PLoS One 8: e74451**

Google Scholar: <u>Author Only</u> <u>Title Only</u> <u>Author and Title</u>

**Smith SR, Gillard JT, Kustka AB, McCrow JP, Badger JH, Zheng H, New AM, Dupont CL, Obata T, Fernie AR, Allen AE (2016) Transcriptional Orchestration of the Global Cellular Response of a Model Pennate Diatom to Diel Light Cycling under Iron Limitation. PLoS Genet 12: e1006490**

Google Scholar: <u>Author Only</u> <u>Title Only</u> <u>Author and Title</u>

**Stecher G, Tamura K, Kumar S (2020) Molecular Evolutionary Genetics Analysis (MEGA) for macOS. Mol Biol Evol 37: 1237-1239**

Google Scholar: <u>Author Only</u> <u>Title Only</u> <u>Author and Title</u>

**Sukenik A, Beardall J, Kromkamp JC, Kopecky J, Masojidek J, van Bergeijk S, Gabai S, Shaham E, Yamshon A (2009) Photosynthetic performance of outdoor Nannochloropsis mass cultures under a wide range of environmental conditions. Aquatic Microbial Ecology 56: 297-308**

Google Scholar: <u>Author Only</u> <u>Title Only</u> <u>Author and Title</u>

**Sukenik A, Carmeli Y, Berner T (1989) Regulation of fatty acid composition by irradiance level in the eustigmatophyte Nannochloropsis sp.1 Journal of Phycology 25: 686-692**

Google Scholar: <u>Author Only</u> <u>Title Only</u> <u>Author and Title</u>

**Takahashi F, Yamagata D, Ishikawa M, Fukamatsu Y, Ogura Y, Kasahara M, Kiyosue T, Kikuyama M, Wada M, Kataoka H (2007) AUREOCHROME, a photoreceptor required for photomorphogenesis in stramenopiles. Proc Natl Acad Sci U S A 104: 19625-19630**

Google Scholar: <u>Author Only</u> <u>Title Only</u> <u>Author and Title</u>

**Takeuchi T, Newton L, Burkhardt A, Mason S, Farre EM (2014) Light and the circadian clock mediate time-specific changes in sensitivity to UV-B stress under light/dark cycles. J Exp Bot 65: 6003-6012**

Google Scholar: <u>Author Only</u> <u>Title Only</u> <u>Author and Title</u>

**Toyooka T, Hisatomi O, Takahashi F, Kataoka H, Terazima M (2011) Photoreactions of aureochrome-1. Biophys J 100: 2801-2809**

Google Scholar: <u>Author Only</u> <u>Title Only</u> <u>Author and Title</u>

**Tragin M, Vaulot D (2018) Green microalgae in marine coastal waters: The Ocean Sampling Day (OSD) dataset. Sci Rep 8: 14020**

Google Scholar: <u>Author Only</u> <u>Title Only</u> <u>Author and Title</u>

**Trapnell C, Roberts A, Goff L, Pertea G, Kim D, Kelley DR, Pimentel H, Salzberg SL, Rinn JL, Pachter L (2012) Differential gene and transcript expression analysis of RNA-seq experiments with TopHat and Cufflinks. Nat Protoc 7: 562-578**

Google Scholar: <u>Author Only</u> <u>Title Only</u> <u>Author and Title</u>

**Vieler A, Wu G, Tsai CH, Bullard B, Cornish AJ, Harvey C, Reca IB, Thornburg C, Achawanantakun R, Buehl CJ, Campbell MS, Cavalier D, Childs KL, Clark TJ, Deshpande R, Erickson E, Armenia Ferguson A, Handee W, Kong Q, Li X, Liu B, Lundback S, Peng C, Roston**

**RL, Sanjaya, Simpson JP, Terbush A, Warakanont J, Zauner S, Farre EM, Hegg EL, Jiang N, Kuo MH, Lu Y, Niyogi KK, Ohlrogge J, Osteryoung KW, Shachar-Hill Y, Sears BB, Sun Y, Takahashi H, Yandell M, Shiu SH, Benning C (2012) Genome, functional gene annotation, and nuclear transformation of the heterokont oleaginous alga Nannochloropsis oceanica CCMP1779. PLoS Genet 8: e1003064**

Google Scholar: <u>Author Only</u> <u>Title Only</u> <u>Author and Title</u>

**Wang D, Ning K, Li J, Hu J, Han D, Wang H, Zeng X, Jing X, Zhou Q, Su X, Chang X, Wang A, Wang W, Jia J, Wei L, Xin Y, Qiao Y, Huang R, Chen J, Han B, Yoon K, Hill RT, Zohar Y, Chen F, Hu Q, Xu J (2014) Nannochloropsis genomes reveal evolution of microalgal oleaginous traits. PLoS Genet 10: e1004094**

Google Scholar: <u>Author Only</u> <u>Title Only</u> <u>Author and Title</u>

**Zones JM, Blaby IK, Merchant SS, Umen JG (2015) High-Resolution Profiling of a Synchronized Diurnal Transcriptome from Chlamydomonas reinhardtii Reveals Continuous Cell and Metabolic Differentiation. Plant Cell 27: 2743-2769**

Google Scholar: <u>Author Only</u> <u>Title Only</u> <u>Author and Title</u>

